# Variant analysis of SARS-CoV-2 strains in Middle Eastern countries

**DOI:** 10.1101/2020.06.18.156810

**Authors:** Khalid Mubarak Bindayna, Shane Crinion

**Affiliations:** Department of Microbiology, Immunology, and Infectious Diseases, College of Medicine and Medical Sciences, Arabian Gulf University, Manama, Bahrain; National University of Ireland Galwa

## Abstract

**Background:** SARS-CoV-2 is diverging from the initial Wuhan serotype, and different variants of the virus are reported. Mapping the variant strains and studying their pattern of evolution will provide better insights into the pandemic spread

**Methods:** Data on different SARS-CoV2 for WHO EMRO countries were obtained from the Chinese National Genomics Data Center (NGDC), Genbank and the Global Initiative on Sharing All Influenza Data (GISAID). Multiple sequence alignments (MSA) was performed to study the evolutionary relationship between the genomes. Variant calling, genome and variant alignment were performed to track the strains in each country. Evolutionary and phylogenetic analysis is used to explore the evolutionary hypothesis.

**Findings:** Of the total 50 samples, 4 samples did not contain any variants. Variant calling identified 379 variants. Earliest strains are found in Iranian samples. Variant alignment indicates Iran samples have a low variant frequency. Saudi Arabia has formed an outgroup. Saudi Arabia, Qatar and Kuwait were the most evolved genomes and are the countries with the highest number of cases per million.

**Interpretation:** Iran was exposed to the virus earlier than other countries in the Eastern Mediterranean Region.

**Funding:** None

## 1. Introduction

Since the discovery in Wuhan in late 2019, the Severe Acute Respiratory Syndrome Coronavirus 2 (SARS-CoV2) virus has spread internationally to 213 countries with confirmed 6,194,533 cases and 376 320 deaths, at time of writing^1^. Of these cases, 536 148 are from the Eastern Mediterranean ^1^. The virus, which causes COVID-19, is a beta-coronavirus related to SARS-CoV and bat coronaviruses^2^. SAR-CoV-2 is easily transmittable due to mutations in the receptor-binding (S1) and fusion (S2) domain of the strain^2^. The rapid spread and high mortality due to the virus have elicited the global response, including the lockdowns and new vaccine development.

While the efforts to understand the virus, dynamics of transmission and epidemiology are still underway, the virus is diverging from the Wuhan strain. The initial Wuhan stain could eventually evolve into a more deadly strain^3^. A recent study by T. Koyama et al.^4^ used 48 genomes to map variants and classify sub-strains from many locations including China, Japan and the USA. We used Koyama’s study model and analysed the strains in the Eastern Mediterranean (EMRO) countries; namely, Iran, Jordan, Kuwait, Saudi Arabia and UAE. The variants were mapped to Wuhan reference genome NC_045512.2 and were aligned using other Wuhan strains. We found variants in 43 of 50 genomes studied.

## 2. Objectives

- To study the genomic and phylogenetic variation in the SARS-CoV2 in the Eastern Mediterranean Region
- To trace the pattern of spread of SARS CoV2 in the Eastern Mediterranean Region

## 3. Methods

### Study Design

Cross-sectional Study

### Sample Size

50

### Source of Data-

We obtained the data from the Chinese National Genomics Data Center (NGDC). Data in the NGDC include those obtained from NGBdb, GenBank, GISAID, GWH and NMDC databases^5^. NCBI Genbank and the Global Initiative on Sharing All Influenza Data (GISAID) were also used to extract data^6,7^. Table 1 shows the source of all 50 samples. 20 of the 50 samples were from Wuhan and used for comparison and validation with the variant analysis performed by Koyama et al.^4^The other 30 samples consisted of 5 samples from Iran, Kuwait, Jordan, Qatar, Saudi Arabia and the United Arab Emirates. These samples were all extracted from GISAID.

**Table 1:**
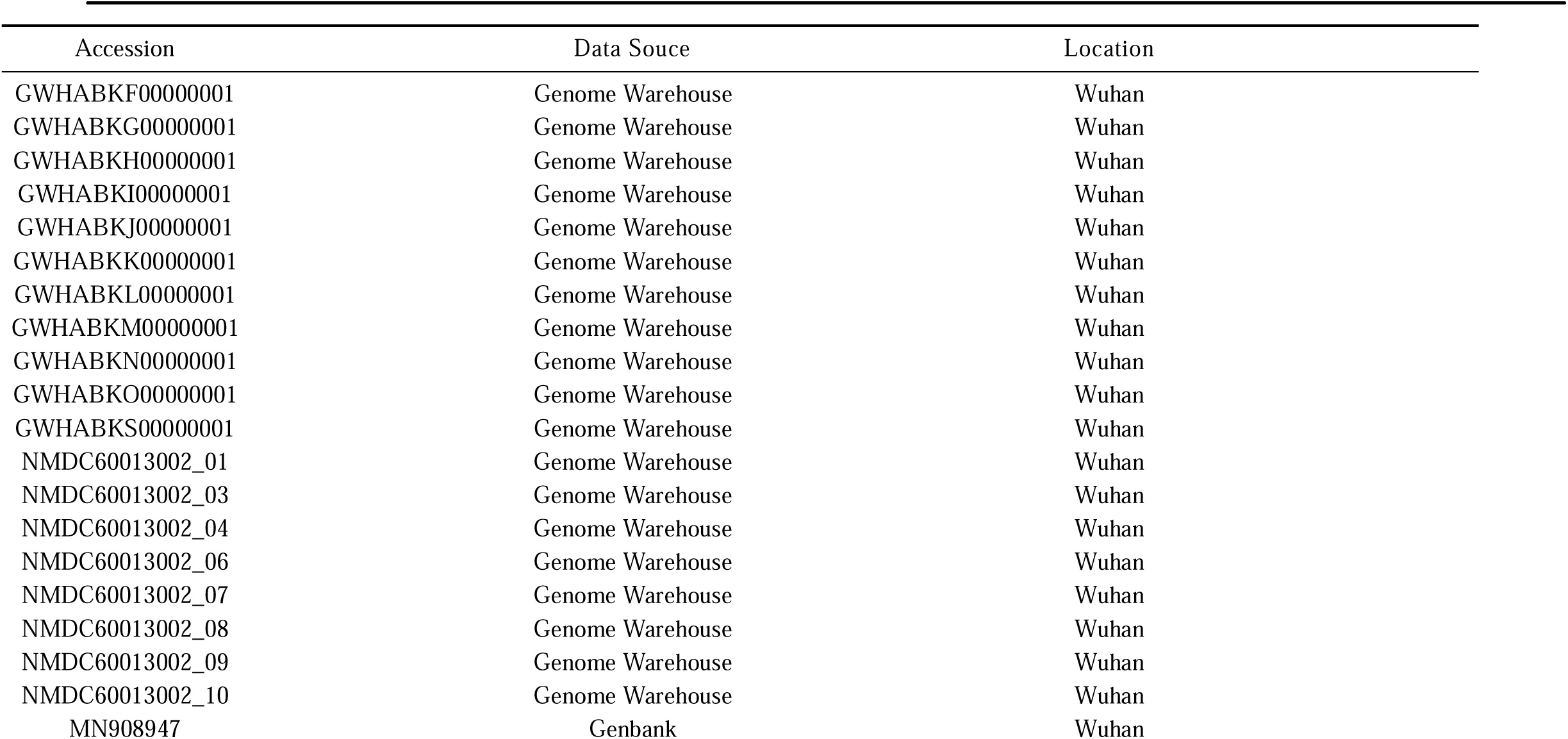
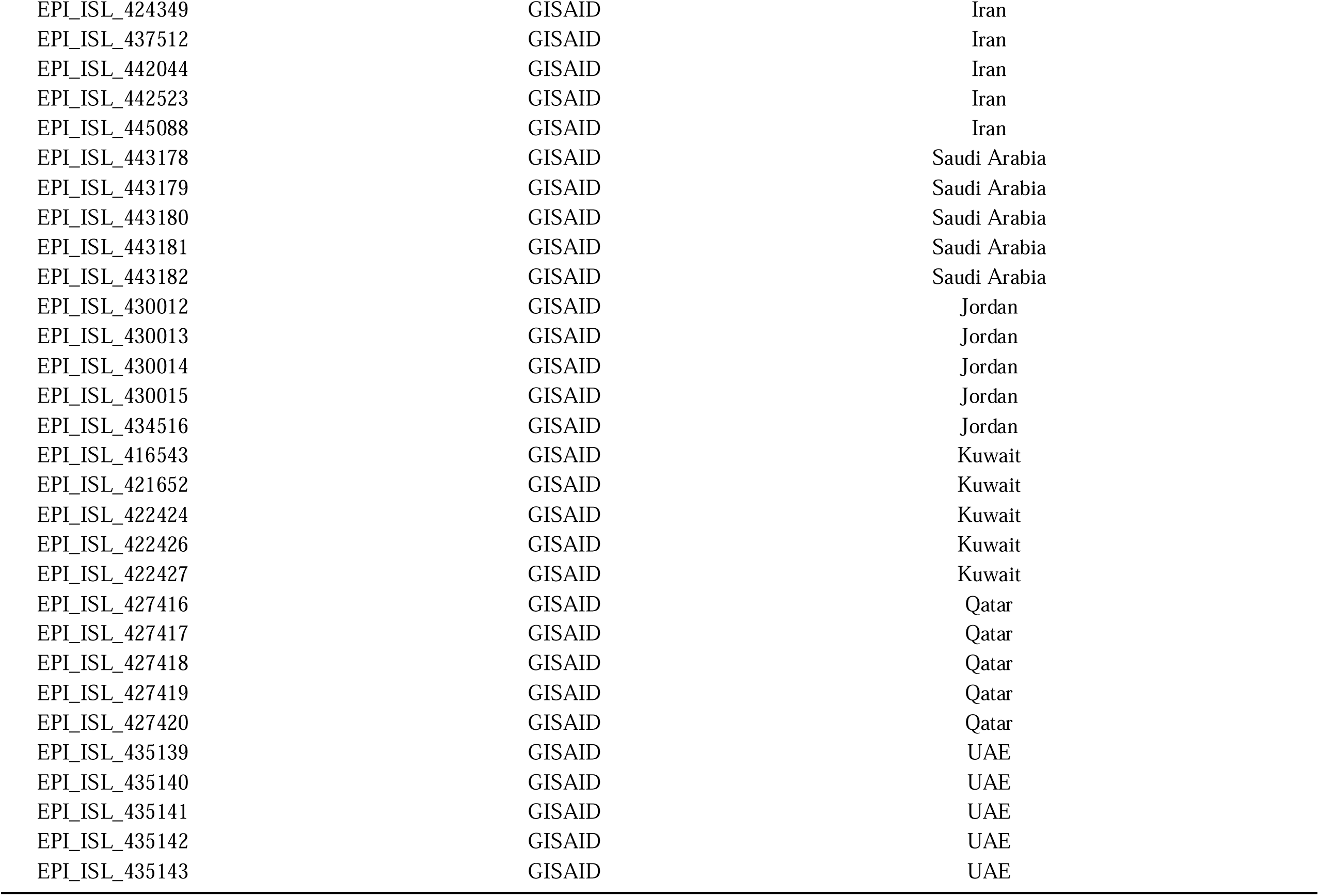
Characterisation of SAR-CoV2 genomes from Middle Eastern populations 5.

### Sample Selection

Samples were selected from WHO EMRO countries with at least 5 high-quality genomes available. Of all the countries, Bahrain, Iraq and Syria were excluded to due to inadequate samples. Lebanon was excluded due to poor quality samples.

### Sample filing

We performed multiple sequence alignments (MSA) using EMBOSS Clustal Omega^8^ and observed for conserved and consensus sequences to study the evolutionary relationship between the genomes. All the outputs were set for the Pearson/FASTA formats. The outputs were represented in sequence alignment file and phylogenetic tree.

We performed variant calling to identify variants that co-occur in different groups and to track the strains in each country. The sequence alignment file from multiple sequence alignments was used to identify the location of variants. The SNP-sites program was used to extract SNP sites from a multi-sample alignment file^9^.

We tabulated the output of the SNP-sites analysis as a variant calling (VCF) file with the list of SNPs against the genotype for each sample.

The SNPs in the VCF file were compared with the SNPs reported by Koyama et al. for the validation.

### Variant annotation

Variant annotation was used to understand the genomic regions and functions affected by the SNPs. We used the Galaxy web platform, and the public server at usegalaxy.org to facilitate variant annotation^10^. SNPeff, a genetic variant annotation program in the Galaxy server, was used to identify the protein level changes caused by SNPs^11^. The genome database for both SARS-CoV-2 NC_045512.2 and SARS NC_004718 were built and compared using SNPeff, and Genbank^7^.

### Genome and variant alignment

We visualised the overlapping variants between populations using the genome alignment. The patterns of the emergence of COVID-19 in each country were analysed. The annotated data was imported, manipulated and plotted using R v3.6.2^12^. dplyr v0.8.4 package was used to summarise and align the data^13^. The visualisation package ggplot2^13^ was used to plot the graphs. The x-axis in the plots indicates the variant position along the SARS-CoV-2 genome; the left y-axis indicates the sample name and the right y-axis represents the country of origin for each sample. This plot is used to compare the genome in different populations.

### Phylogenetic analysis

Evolutionary and phylogenetic analysis is used to explore the evolutionary hypothesis for the strain emergence in each country. Bayesian Evolutionary Analysis Sample Trees (BEAST) v1.10.4, is used to perform Bayesian analysis of molecular sequences using MCMC^14^. The alignment file output from MSA was used as the input data for Bayesian analysis.

The HKY transition-transversion parameter, a burn-in of 1×10^6^ iterations and a Coalescent tree were used as models for molecular evolution here. The unrooted tree obtained from the models and the phylogenetic tree generated by Clustal Omega is used to predict the associations between samples.

## 4. Results

Of the initial 50 samples, 20 of them were from Wuhan, 4 of them did not contain any variants (GWHABKI00000001, GWHABKL00000001, NMDC60013002_08 and MN908947), The SARS-CoV-2 had the best scoring variant annotation (Table 2). The results from the variant annotation are presented in Figure 1. However, ten variants caused errors and are not included in the figure. Multi-allelic variants were included in Figure 1 if the second alternative allele was likely due to poor reading.

**Table 2.**
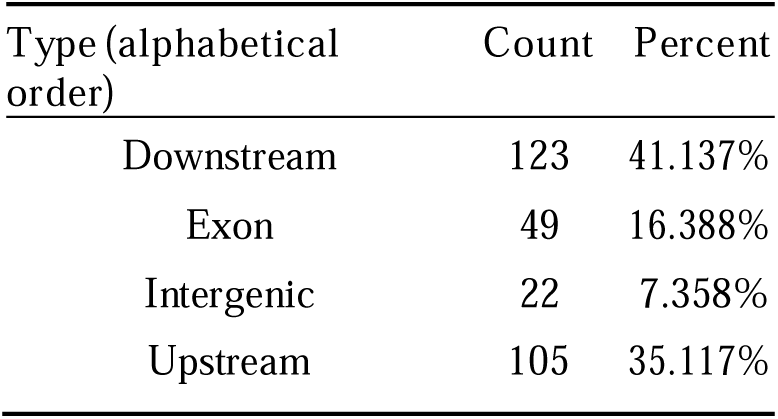
The number of effects by region of the variant.

**Fig. 1.**
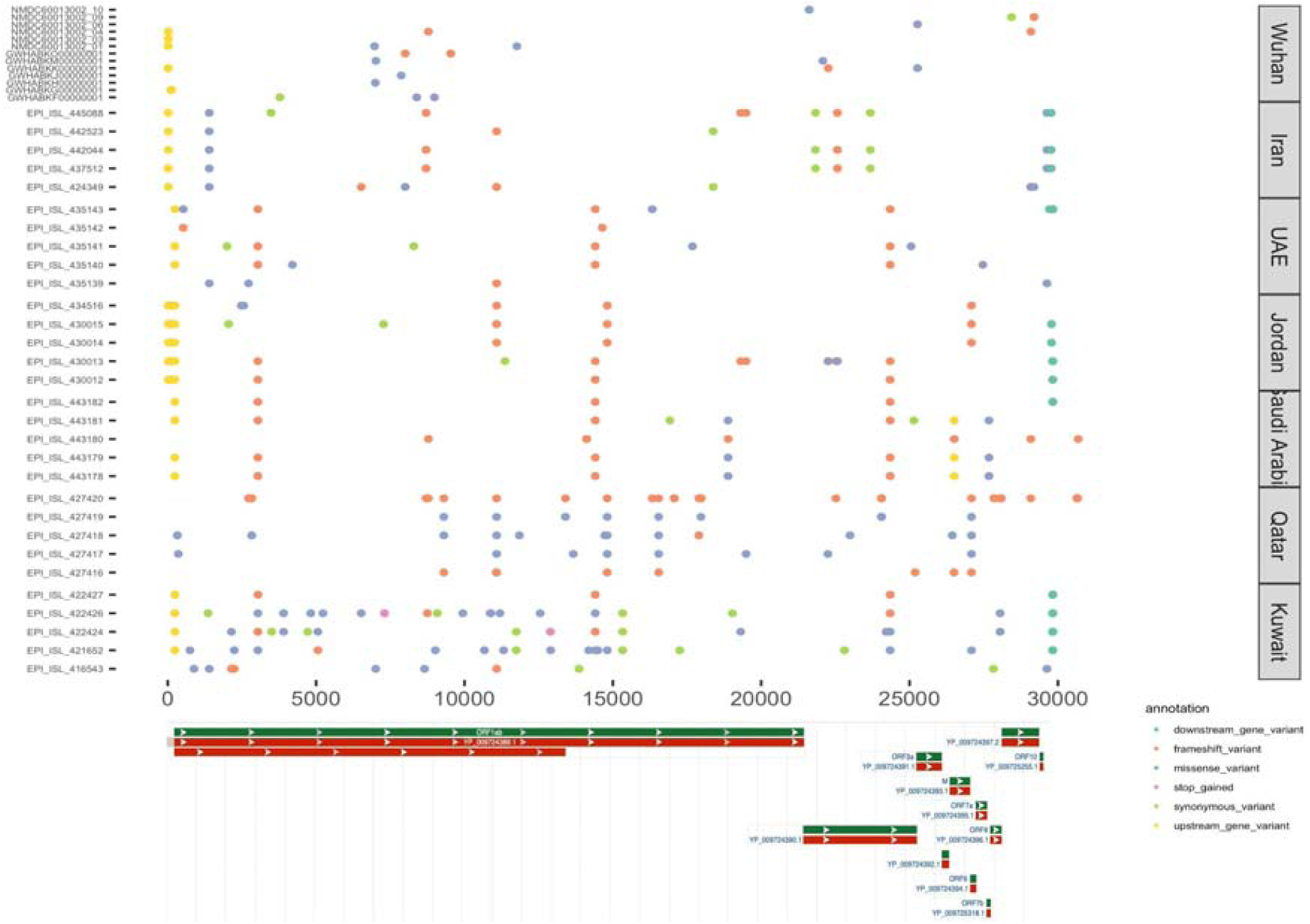
A graphical representation of the variants found in COVID-19 genomes. Samples are split by country of origin. The gene structure was extracted from NCBI Genome Browser for Wuhan reference genome NC_045512.2. Graphical representation was generated using R and ggplot2.

The distribution of SNPs across samples (studied using the SNP genotypes from 442 SARS-CoV-2 strain) showed a vast difference between the sub-strains within each country. Variant calling identified 379 variants. Of these, 250 were modifier variants, 21 were modifier variants, 18 were low impact variants, and 10 were high impact variants. The variants had a missense/silent ration of 1.75. Table 2 shows the distribution of variants by region, and Table 3 shows the distribution of variants by their type. Figure 3 shows the number of transitions and transversions observed in the sample. Figure 2 shows the drift towards transition is evident with a Ts/Tv ratio of 3.36. One variant occurs for every 378 bases and 16 multi-allelic sites (all include indels).

**Table 3.**
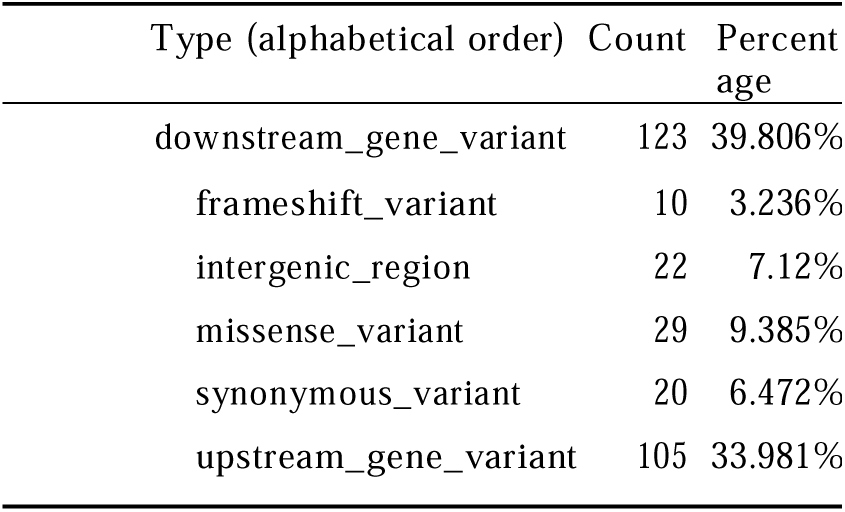
The number of effects by type of the variant.

**Figure 2.**
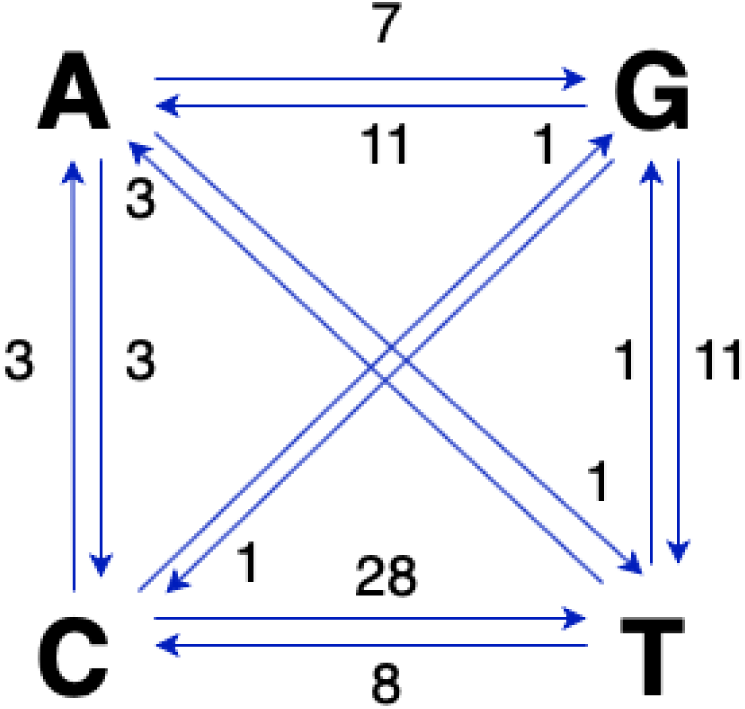
Distinct base-pair changes among the SARS-CoV-2 genome.

**Figure 3:**
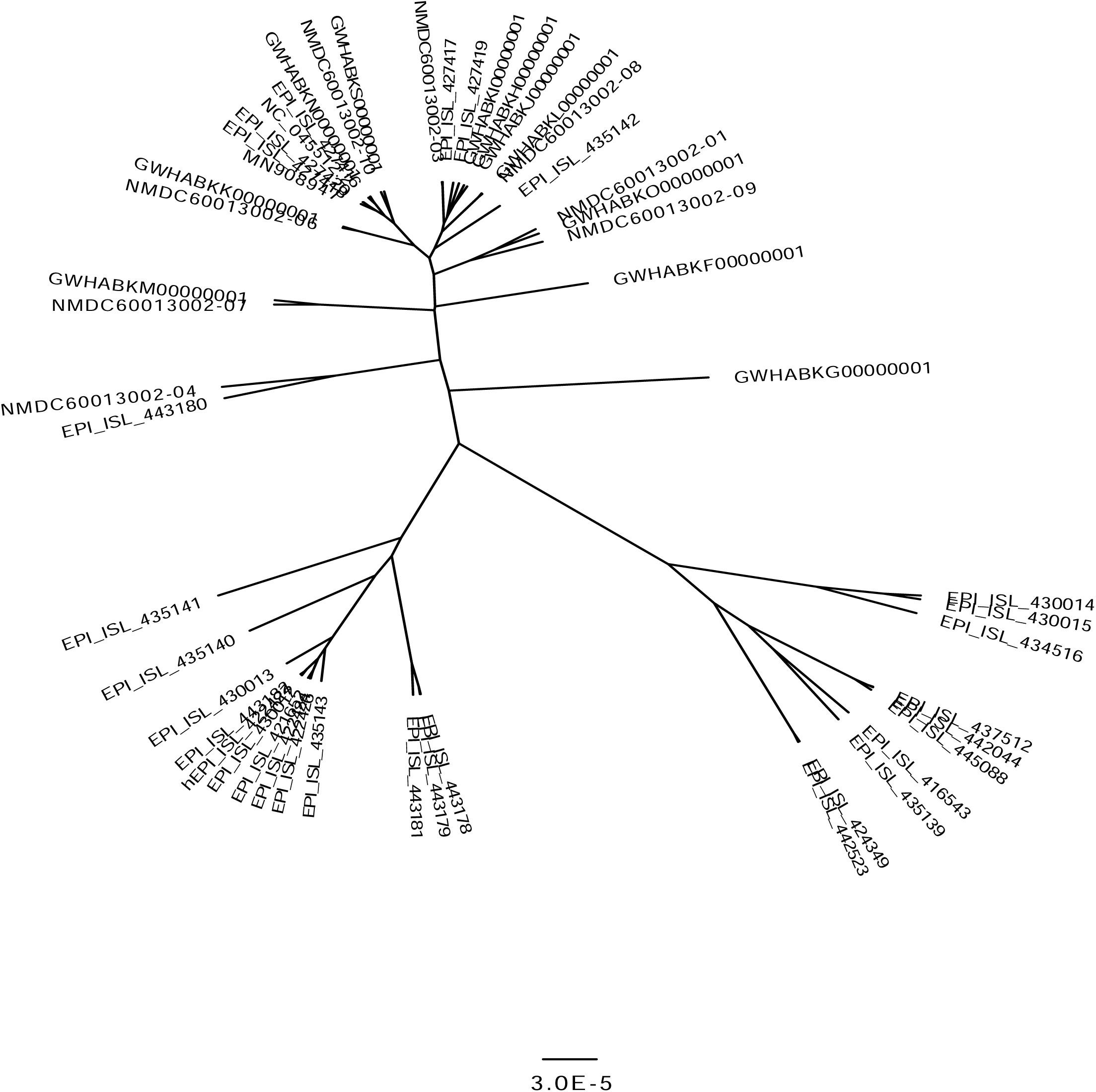
Consensus sequence from BEAST analysis.

From the phylogenetic analysis and tree generation in Figure 3, we can infer that the Iranian samples have the earliest common ancestor. Saudi Arabia samples are all form a distinct group compared to other samples. The samples from SA, Qatar and Kuwait have the most developed SARS-CoV2 genomes.

## 5. Discussion

Four of the Wuhan strains that did not have any variants matched with the variants identified by Koyama et al. This indicates that the present study is valid.

### Variant-based assessment

We used MSA results to create a dot-plot and to visualise the variants in the sample. The variants found in different regions, in the descending order of their number were from Wuhan, Iran, UAE, Jordan, Saudi Arabia, Qatar and Kuwait. The proximity of Iranian strains to the Wuhan strain is not surprising as the first recorded cases in Iran were the individuals travelling back from China.

When analysed for the distribution of infections per capita, Qatar (19,211 per 1 million) stands at the top, followed by Bahrain (6,335) and Kuwait (6,142). No sample from Bahrain was available when this study was conducted. However, Qatar and Kuwait show a significant number of variants. These variants might be associated with the higher symptomatic cases in these regions compared to the others. These countries have small populations (< 5 million), and faster genetic drift in these countries is expected.

Many of the initial cases in Bahrain had travelled from Iran. The alignment observed in the present study also allowed us to track the path of transmission from Wuhan. Although the United Arab Emirates reported the first confirmed case of corona infection in the middle east^15^, the first infection might have occurred in Iran, from which it eventually spread across the middle east.

### Phylogenetic-based approach

Phylogenetic trees help in understanding the evolutionary relationships between groups. In the present context, they are used to identify the earliest strains and to track the spread of COVID-19 across the middle east. We expected strains similar to Wuhan to have emerged earlier than the other strains.

Saudi Arabia has reported the highest number of cases in the Middle East (WHO, 2020). This can be explained from our phylogenetic analysis. The four samples from Saudi Arabia (EPI_ISL_443181, EPI_ISL_443180, EPI_ISL_443179, EPL_ISL_443178) were more distantly related than samples from any other country. This distinction might be due to the swift response by the country leading to a unique and restricted strain. The remaining Saudi Arabian samples (EPI_ISL_443182) were in a separate clade with one sample from Wuhan (GWHABKM00000001), one from Kuwait (EPI_ISL_422426), and one from UAE (EPI_ISL_435143).

Jordan has adopted a stringent pandemic response strategy. They imposed restriction and closed the non-essential services early on. The same is reflected in the genome variation. Four out of the five Jordan samples (EPI_ISL_430015, EPI_ISL_430014, EPI_ISL_430013, EPI_ISL_430012) were in a separate clade with only one sample (MN908947) similar to Wuhan. The Jordan samples clusters at one of the final tree branching, indicating that they are the most evolved variants. Their sample divergence might be the reason for their most distinctive genome.

The remaining Jordan sample (EPI_ISL_434516) and a Qatar sample (EPI_ISL_427419) diverge from a Kuwait sample (EPI_ISL_421652), and all of these have emerged from the common Wuhan ancestor (GWHABKJ00000001)

One of the earliest lineage divergences are found in five samples from Iran (EPI_ISL_445088, EPI_ISL_442044, EPI_ISL_442523, EPI_ISL_437512, EPI_ISL_424349), two from Wuhan (GWHABKN0000001, NMDC60013002-07), one from UAE (43519) and one from Kuwait (EPI_ISL_416543). It strongly reiterates our hypothesis that the Iranian’s introduced COVID-19 to other middle eastern countries. The only branchings that precede the cluster of Iranian samples are four samples from Wuhan (NC_044512.2, NMDC60013002-04, NMDC60013002-08, GWHABKS00000001) and one sample from the UAE (EPI_ISL_435141). The UAE reported the first case of COVID-19 in the middle east^15^.

## 6. Conclusion

Here we trace the spread of COVID-19 using variant and phylogenetic analysis. This study reveals the structure of spread among populations. We conclude that the earliest strains are found in Iranian samples. Iran was exposed to the virus earlier than in other countries.

Kuwait and Qatar have a high frequency of novel variants due to small populations size, which leads to an accumulation of mutations. As Bahrain also has the highest number of infections per million, it is expected that Bahranian genomes would also have a high variant rate. As all the samples from Saudi Arabia are experiencing some differentiation and Saudi Arabia reports a high death rate, more vigilance is necessary to prevent these outgroups from contributing to a more severe sub-strain with higher mortality rate.

## Acknowledgements

The authors are grateful for the timely sequencing and release of genomes to make this study possible.and for Dr. Anusha C P for her comments.

## Funding

This research received no specific grant from any funding agency in the public, commercial, or not-for-profit sectors.

## Notes

### Competing Interest Statement

The authors have declared no competing interest.

